# Effects of surface geometry on light exposure, photoacclimation and photosynthetic energy acquisition in zooxanthellate corals

**DOI:** 10.1101/2023.07.16.549221

**Authors:** Tomás López-Londoño, Susana Enríquez, Roberto Iglesias-Prieto

## Abstract

Symbiotic corals display a great array of morphologies, each of which has unique effects on light interception and the photosynthetic performance of *in hospite* zooxanthellae. Changes in light availability elicit photoacclimation responses to optimize the energy balances in primary producers, extensively documented for corals exposed to contrasting light regimes along depth gradients. Yet, response variation driven by coral colony geometry and its energetic implications on colonies with contrasting morphologies remain largely unknown. In this study, we assessed the effect of the inclination angle of coral surface on light availability, short- and long-term photoacclimation responses, and potential photosynthetic usable energy. Increasing surface inclination angle resulted in an order of magnitude reduction of light availability, following a linear relationship explained by the cosine law and relative changes in the direct and diffuse components of irradiance. The light gradient induced by surface geometry triggered photoacclimation responses comparable to those observed along depth gradients: changes in the quantum yield of photosystem II, photosynthetic parameters, and optical properties and pigmentation of the coral tissue. Differences in light availability and photoacclimation driven by surface inclination led to contrasting energetic performance. Horizontally and vertically oriented coral surfaces experienced the largest reductions in photosynthetic usable energy as a result of excessive irradiance and light-limiting conditions, respectively. This pattern is predicted to change with depth or local water optical properties. Our study concludes that colony geometry plays an essential role in shaping the energy balance and determining the light niche of zooxanthellate corals.

## Introduction

The combination of the symbioses with photosynthetic dinoflagellates and colonial integration is thought to be a key factor for the ecological and evolutionary success of corals in tropical coral reef ecosystems [1,2]. The nutritional symbiosis allows this group of sessile benthic animals to efficiently use solar radiation as major source of energy, conferring important metabolic advantages in the predominantly oligotrophic environments in which they evolved [3,4]. On the other hand, colony integration allows resource translocation from source to sink sites according to metabolic needs [5–7], contributing to their competitive dominance over other benthic invertebrates by strengthening colony-wide fitness and stress resistance [8,9].

Colony integration in corals led to the evolution of a complex array of morphologies under the influence of environmental factors [10,11]. Changes in colony morphology and surface geometry have important implications on the light interception capacity [12,13] and the consequent photosynthetic energy acquisition by the whole colony [14,15]. The light interception at a particular colony surface area is partially mediated by the cosine of the angle between the direction of incident irradiance and that particular surface, a phenomenon based on geometrical relations usually referred as the Lambert’s Cosine Law [12,16]. This law states that there is an angle dependent reduction in the amount of light energy intercepted by a surface, following a cosine function. Such physical law may explain why coral species tend to adopt horizontal plate-like morphologies in light-limited environments, such as caves and deep-water, in order to maximize light interception, and branching or foliaceous morphologies in high-light environments to facilitate light attenuation and reduce the coral surface area exposed to intense irradiance and thus the potential of photoinhibition [12,14,15].

Contrasting light climates elicit diverse photoacclimation responses in zooxanthellate corals, as in all primary producers, to minimize the imbalance between the absorbed energy and the energy that can be incorporated through photochemistry or dissipated by photoprotective mechanisms [17]. Readjustments of the photosynthetic apparatus of the zooxanthellae occur over both short- and long- term time scales. In the short-term, the energy imbalance can lead to, e.g., reductions in the energy transfer efficiency to the photosystem II, which is the protein complex responsible for the release of oxygen and electrons from photosynthetic water oxidation. In the long-term, the energy imbalance can lead to changes in the density of light-harvesting pigments and/or electron-consuming sinks, affecting the photosynthetic electron flow capacity [17,18]. Extensive documentation exists regarding the change in photoacclimatory responses of corals exposed to environmental gradients, primarily focusing on the variation associated with bathymetric differences and species-specific physiological capabilities of symbiotic dinoflagellates [19–24]. Recent research has also highlighted significant variation of photoacclimatory responses associated with seasonal changes in light and temperature [25]. The use of non-destructive methods to measure some of these photoacclimatory responses, such as active chlorophyll *a* fluorescence induction, has gained widespread appeal in recent decades, making it possible to obtain reliable data on photoacclimation processes while preserving the integrity of coral colonies [23].

The amount of light that reaches coral colonies directly affects the photosynthetic capacity of *in hospite* zooxanthellae, which in turn determines the photosynthetic usable energy that can be translocated to their host [26]. Both high and low light conditions can potentially limit the translocation of usable energy and impact the overall energetic performance of coral holobionts. In high-light environments, the photosynthetic energy reduction occurs due to the increased costs of metabolic maintenance in the zooxanthellae when photosynthesis is fully saturated. Conversely, in low-light environments it occurs as a consequence of the reduction in the photosynthetic activity and photosynthetically fixed energy [26]. Considering coral colonies with complex morphologies exposed to heterogeneous surface light levels, it can be predicted that there are gradients of photosynthetic productivity within the colony. Areas at with a particular angle of inclination may provide maximum outputs of photosynthetic energy, while other areas may exhibit minimum productivity or even negative carbon balances. Coral surfaces with high productivity can serve as a source of metabolic energy for other portions of the colony that are exposed to excessive irradiance and thus increased metabolic maintenance expenditure (e.g., horizontal surfaces exposed to direct sunlight), or to areas under light-limited conditions where there is very low fixed energy income through symbiont photosynthesis (e.g., vertical, self-shaded surfaces). These complex interactions between light availability, colony morphology, and photosynthetic productivity highlight the intricate energy dynamics within coral colonies.

The aim of this study was to investigate the influence of the inclination angle of coral colony surfaces on light exposure, short- and long-term photoacclimatory responses, and the acquisition potential of photosynthetic usable energy. For this purpose, we conducted an *in-situ* experiment with small samples of the coral species *Orbicella faveolata* exposed to different inclination angles.

Although this species can form colonies with massive and complex morphologies, the predominant flat morphology at the meso-scale limits the formation of significant surface light gradients, making it an ideal system to study local photoacclimation associated with coral surface inclination. Our findings revealed substantial variations in light availability and photoacclimation responses resulting from changes in coral surface inclination, similar to those commonly observed along depth gradients. Furthermore, we documented here the impact of surface inclination on local productivity, energetic demands and energy fluxes within coral colonies.

## Materials and Methods

### Sampling design

Twenty coral fragments of the species *Orbicella faveolata* of similar size (∼5x5cm) formed the basis for this experimental study. Experimental coral fragments were salvaged from projects conducted at the Reef System Unit in Puerto Morelos, Mexico, and were lying in a flat, horizontal surface at a constant depth of 5 m since at least one year before experiments. Experimental corals were glued with underwater epoxy (Z-Spar A-788 epoxy) to PVC couplers and fixed to a panel at 3 m depth over a seagrass bed to naturally reduce the upwelling irradiance. Coral samples were initially acclimated for two weeks at an intermediate angle of exposure to downwelling irradiance of 45°, and then evenly distributed across five inclination settings facing north: 0° (horizontal), 25°, 45°, 65° and 90° (vertical). Massive coral colonies in their natural habitat at 5 m depth were also used for some analyses. Photoacclimatory responses of both experimental corals and colonies in their natural habitat, were assessed by using non-destructive techniques only.

### Quantum yield of charge separation in PSII

Chlorophyll *a* (Chl *a*) fluorescence was recorded daily in all coral fragments (N = 20, n = 4 per inclination setting) after adjusting the inclination settings until perceiving a steady state photochemical activity of photosystem II (PSII). The maximum (*F*_v_/*F*_m_) and effective (Δ*F*/*F*_m_’) quantum yields of PSII were respectively recorded at dusk and at local noon using a submersible pulse amplitude modulated fluorometer (Diving-PAM, Walz, Germany) (saturation light pulse width 0.6s of >4500µmol quanta m^-2^ s^-1^). Measurements of *F*_v_/*F*_m_ and Δ*F*/*F*_m_’ were also recorded on coral surfaces along an inclination gradient of three massive *O. faveolata* colonies in their natural habitat at a constant depth of approximately 5 m. *In situ* data were collected along transects facing north laid from top to bottom of the colonies, controlling the inclination angle with a buoyancy device attached to a customized protractor. The maximum excitation pressure over PSII (*Q*_m_) was calculated as *Q*_m_ = 1 - [(Δ*F*/*F*_m_’ at noon) / (*F*_v_/*F*_m_ at dusk)] [27]. When measuring Δ*F*/*F*_m_’ on the experimental fragments, total irradiance was also recorded at every inclination setting using the cosine-corrected PAR sensor of the Diving-PAM, previously calibrated against a reference quantum sensor (LI- 1400; LI-COR, USA). The relative variation in the direct and diffuse components of irradiance was determined on a single day at noon, by shading the PAR sensor with a black panel to remove the direct component. The diurnal variation of the quantum yield of PSII in the experimental corals was measured over a diurnal cycle (06:00-19:00) on a cloudless day at the end of the photoacclimatory period.

### Optical properties of coral tissues

The light absorption capacity of intact corals was determined spectrophotometrically following the technique initially developed by Shibata (28) and subsequently modified [29,30]. The reflectance (*R*) spectra of all experimental corals (N = 20, n = 4 per inclination setting) were measured 10 months after adjusting the inclination settings, using a modular spectrophotometer (Flame-T-UV-VIS, Ocean Optics Inc.). The fraction of incident light absorbed by the coral tissue, absorptance (*A*), was calculated as *A* = 1 - *R*, assuming that the light transmitted through the coral skeleton is negligible [29]. The absorptance peak of chlorophyll *a* (Chl *a*) at 675 nm (*A*_675_) was used as a proxy of relative changes in Chl *a* content, considering that light absorption at this wavelength has minimal interference from accessory algal and animal pigments [29]. The minimum quantum requirement of photosynthesis (1/*Φ*) was calculated based on the amount of light being absorbed and used to drive photosynthesis in the linear increase of the P vs E curve at sub-saturating irradiance (see below). The ratio of absorbed light was calculated by multiplying the light emission spectrum of the lamp used in the P vs E curves by the *in vivo* absorption spectrum of corals [31].

### Photosynthetic parameters

Photosynthetic parameters derived from P vs E (photosynthesis vs irradiance) curves were obtained from 15 experimental corals, 12 months after exposing the samples to the experimental conditions. For the analysis, three coral samples from each inclination setting that maintained predominantly flat surfaces were selected, avoiding the formation of internal light and productivity gradients during measurements. Photosynthesis and respiration rates were measured with a fiber- optic oxygen meter system (FireSting, Pyroscience) in a custom-made acrylic chamber with filtered seawater (0.45 µm) under constant agitation. Water temperature was maintained at 28 °C using an external circulating water bath (Isotemp, Fisher Scientific, USA). Ten levels of irradiance (0, 37, 65, 97, 147, 232, 361, 479, 677 and 904 µmol quanta m^-2^ s^-1^) were supplied at 10 min intervals with a dimmable LED (Philips, China) and a combination of neutral density filters. Photosynthetic parameters were calculated as in [31], normalized to coral surface area estimated with the aluminum foil technique [32]. Briefly, the respiration rates were calculated by averaging the dark respiration rates obtained before illumination (*R*_D_) and after exposing the corals to the maximum level of irradiance (R_L_). Maximum photosynthetic rates (*P*_max_) were calculated using the averaged values of photosynthesis obtained above the saturating irradiance (*E*_k_). The compensating irradiance (*E*_c_) corresponded to the light intensity at which the rate photosynthesis matched the rate of respiration. The slope of the linear increase of photosynthesis at sub-saturating irradiance (or photosynthetic efficiency, *α*), was calculated using least-square regression analyses. This last parameter was also used for the calculation of 1/*<λ*.

### Local irradiance regimes

Data of sea surface irradiance (*E*_0_) recorded by a nearby (∼100 m) meteorological station (Estación Meteorológica Puerto Morelos) were used to characterize the temporal variation of solar energy and light-driven processes according to the inclination angle of coral samples (n = 143 days). Total downwelling irradiance at the experimental depth (*E*_z_) was calculated following the Lambert-Beer law (*E*_z_ = *E*_0_ e^-*K*d^ ^z^) [16], using a vertical attenuation coefficient for downwelling irradiance (*K*_d_) of 0.20 m^-1^ [33]. Light availability at each inclination setting (*E*Σ) was estimated using correction factors derived from empirical measurements of local irradiance at noon (1.00 at 0°, 0.85 at 25°, 0.58 at 45°, 0.34 at 65°, and 0.09 at 90°). The proportion of daily light integral (DLI) corresponding to sub-compensating (below *E*_c_), compensating (between *E*_c_ and *E*_k_), and saturating light intensities (above *E*_k_) were also calculated for each inclination setting. The daytime period during which the light intensity exceeded the compensating irradiance (*H*_com_) and saturating irradiance (*H*_sat_) were calculated as in Dennison and Alberte (34). *H*_com_ was used as descriptor of the daytime period (h) during which irradiance was enough for photosynthesis to compensate the respiratory demands of the coral holobiont, while *H*_sat_ was used as descriptor of the amount of time during which irradiance exceeded the maximum photosynthetic capacity of corals.

### Light-driven processes

Based on the availability of light and the photosynthetic parameters derived from the P vs E curves, we estimated the relative change in coral productivity for each inclination setting. The photosynthetic productivity (*P*_g_) over diurnal cycles was estimated following Jassby and Platt (35) as: *P*_g_ = *P*_max_ tanh (*α E*Σ/ *P*_max_). We assumed that not all algal photosynthates are translocated to their coral hosts, considering the energy expenditure of the zooxanthellae associated with repairing its photosynthetic apparatus from photodamage which is proportional to light availability [26]. The amount of photosynthetic usable energy supplied (*PUES*) was calculated by subtracting the estimated costs of repair (*C*_a_) from the photosynthetic output of the zooxanthellae (*P*_g_) [26].

### Data analysis

Simple linear regression models were used to explore the explanatory power of coral surface inclination angle on the variation of local irradiance and coral physiology and optics. An analysis of covariance was conducted to test for differences in the linear regression describing the association of photosynthetic parameters with coral surface inclination in experimental samples and intact colonies in their natural habitat (interaction of parameters with inclination angle of coral surfaces). Pearson- product correlation analysis were conducted to explore the type and magnitude of the relationship between some variables (respiration vs photosynthetic rates, and maximum quantum yield of PSII vs minimum quantum requirement). Analyses were conducted using R version 4.2.2 [36].

## Results

Coral samples of *Orbicella faveolata* were exposed to contrasting light climates by adjusting the inclination angle of the coral surface, which spanned from 0° (fully horizontal orientation) to 90° (fully vertical orientation). Maximum irradiance recorded at local noon available for horizontally oriented corals was 735.67 ± 68.67 (mean ± standard deviation) μmol quanta·m^-2^·s^-1^, while for vertically oriented corals it was 71.04 ± 8.06 μmol quanta·m^-2^·s^-1^. In relative units, the available irradiance for corals at 0° and 90° inclination angles was estimated as 55% and 5% of the incident irradiance at sea surface, respectively. A linear regression analysis revealed that surface inclination angle explained 95% of the variation in maximum irradiance available for the experimental corals (**Fig 1A**; R^2^ = 0.95, p < 0.001). The direct and diffuse components of local irradiance exhibited opposite patterns with respect to coral surface inclination (**Fig 1B**). In vertically oriented corals, the diffuse component accounted for most of the available irradiance (81%), while for horizontally oriented corals, it was the direct component that contributed the most to the available irradiance (86%). Furthermore, a strong positive correlation was found between the direct component of local irradiance and the cosine of the angle of coral surface (r_(3)_ = 0.99, p < 0.001), confirming that coral surfaces behave as a cosine-corrected light collectors at this scale [29], in accordance with Lambertian cosine law [16].

**Fig 1.**
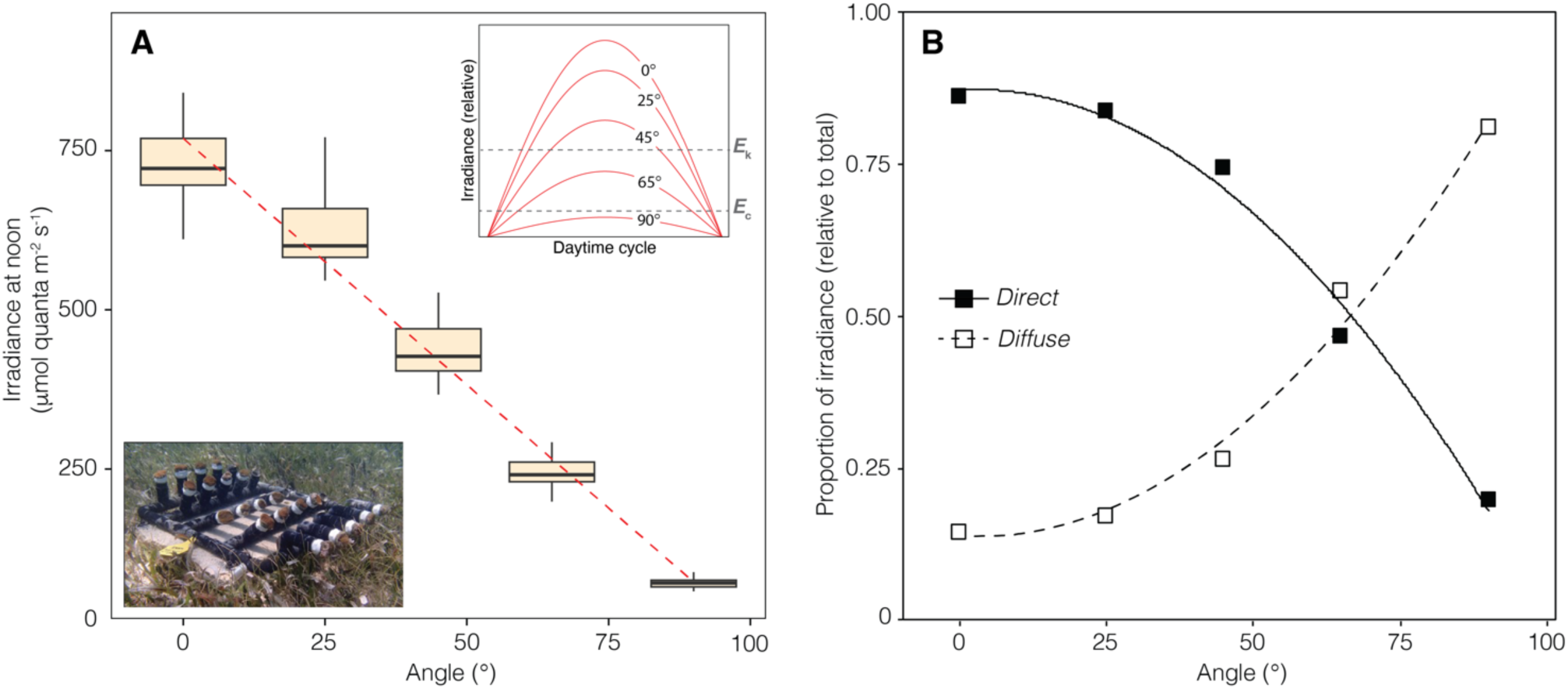
Variation in the local light climate with the inclination angle of coral surface. (A) Box plots of the peak irradiance at local noon with a linear regression describing the relationship between irradiance and surface inclination. The top right insert depicts the predicted pattern of diurnal variation of irradiance for each inclination setting, indicating the relative positions of the compensating irradiance (*E*c) and saturating irradiance (*E*k). The bottom-left insert shows the PVC structure used in the experiment to adjust the inclination angle of coral samples. (B) Relative variation of the diffuse and direct components of total irradiance. Cosine functions were used to illustrate the relationship between the direct and diffuse components of irradiance with surface inclination.

### Quantum yield of charge separation in PSII

After nearly five days of acclimation to the inclination gradient, the maximum excitation pressure over photosystem II (PSII) at noon (*Q*_m_) was found to be more than five times higher in horizontally oriented corals (0.56 ± 0.08) than in vertically oriented corals (0.10 ± 0.04) (**Fig 2A**). Absolute values of the maximum (*Fv/Fm*) and effective (τι*F/Fm*’) quantum yields of PSII over the diel cycle showed a gradual reduction of the PSII photochemical activity after dawn, reaching minimum *ΔF/Fm*’ values at midday and then gradually recovering at dusk until reaching former dawn *Fv/Fm* levels in all inclination settings. However, important differences were observed in the maximum reductions of Δ*F*/*F*_m_’ at noon relative to maximum *F*_v_/*F*_m_ values at dusk/dawn along the inclination gradient, being much more pronounced in horizontally oriented corals (average reduction of 62%) compared to vertically oriented corals (average reduction of 13%) (**Fig 2B**, see **Supplementary Table** for all parameters estimates). The greater reduction of Δ*F*/*F*_m_’ relative to *F*_v_/*F*_m_ in horizontal or slightly inclined coral samples reflects a larger induction of non-photochemical quenching (NPQ), a photoprotective mechanism to dissipate excess absorbed light energy as heat. The complete recovery of *F*_v_/*F*_m_ at the end of the day relative to former dawn levels at each inclination setting suggests full relaxation of NPQ, supporting the notion that the corals had already undergone stable photoacclimation to the local light climates at each inclination setting.

**Fig 2.**
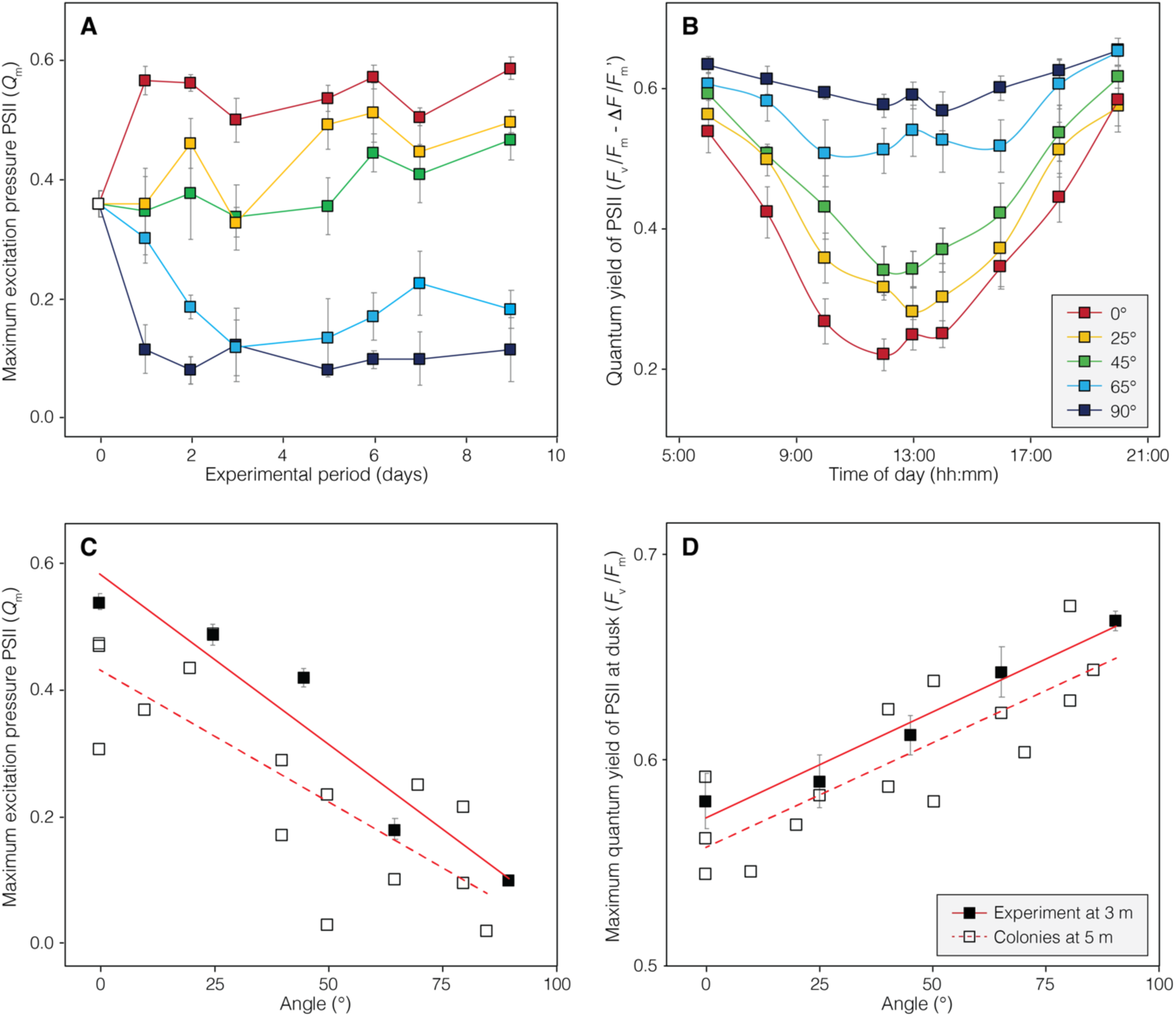
Variation in chlorophyll *a* fluorescence parameters with coral surface inclination. (A) Maximum excitation pressure over PSII (*Q*m) since the first day of exposure to the experimental conditions until it reached an apparent steady state. (B) Diurnal oscillation of the maximum (*Fv/Fm*) and effective (*ΔF/Fm’*) quantum yield of the PSII. The linear associations of *Q*m and *Fv/Fm* with coral surface inclination are shown in plots (C) and (D) respectively, for the experimental samples at 3 m depth (solid squares and continuous lines) and field measurements on coral colonies in their natural habitat at 5 m depth (empty squares and discontinuous lines). Error bars in all graphs are SE.

Significant linear associations between all photosynthetic descriptors derived from chlorophyll *a* (Chl *a*) fluorescence measurements and coral surface inclination angle were also found, both for colonies under natural environmental conditions and for the experimental samples. Linear regression analyses indicated that the inclination angle of coral surface explained 78% of the *Q*_m_ variation in experimental corals at 3 m depth and 63% of the within-colony variation in natural environments at 5 m depth (R^2^ = 0.78, p < 0.001 and R^2^ = 0.63, p < 0.001, respectively) (**Fig 2C**). Experimental corals were located at 3 m depth, which explains their slightly higher *Q*_m_ values compared to those of natural colonies at 5 m depth. This, as *Qm* quantifies the maximum excitation pressure on PSII at noon, expected to be greater in corals exposed to higher-light conditions. Coral surface inclination also explained 68% of the within-colony variation of *F*_v_/*F*_m_ in natural environments (R^2^ = 0.68, p < 0.001) and 36% of the *F*_v_/*F*_m_ variation in experimental samples (R^2^ = 0.36, p < 0.001) (**Fig 2D**). The covariance analysis (condition by angle interaction) indicated that the pattern of change in both *Q*_m_ and *F*_v_/*F*_m_ as a function of coral surface inclination was similar in natural environments and experimental conditions (*F*_(1,91)_ = 1.95, p = 0.17 and *F*_(1,91)_ = 0.001, p = 0.97, respectively).

### Optical properties of coral tissues

Variation in coral absorptance at 675 nm (*A*_675_), which corresponds to the peak absorption wavelength of chlorophyll *a* (Chl *a*), was employed as a non-destructive proxy to assess changes in Chl *a* content in the experimental corals. This proxy indicated that coral pigmentation also varied as a function of the inclination angle of coral surfaces. Linear regression analysis revealed that the inclination angle accounted for only 21% of the variation in *A*_675_, with a marginal level of significance (R^2^ = 0.21, p = 0.042). The greatest reductions in *A*_675_ were observed in horizontal and slightly inclined coral samples (0.83 ± 0.06 at 0° and 0.84 ± 0.07 at 25°) indicating a decrease in pigment content. In contrast, the highest values were observed in corals at intermediate and greater inclination angles (0.90 ± 0.02 at 45° and 90°) suggesting maximum pigment content (**Fig 3A**).

**Fig 3.**
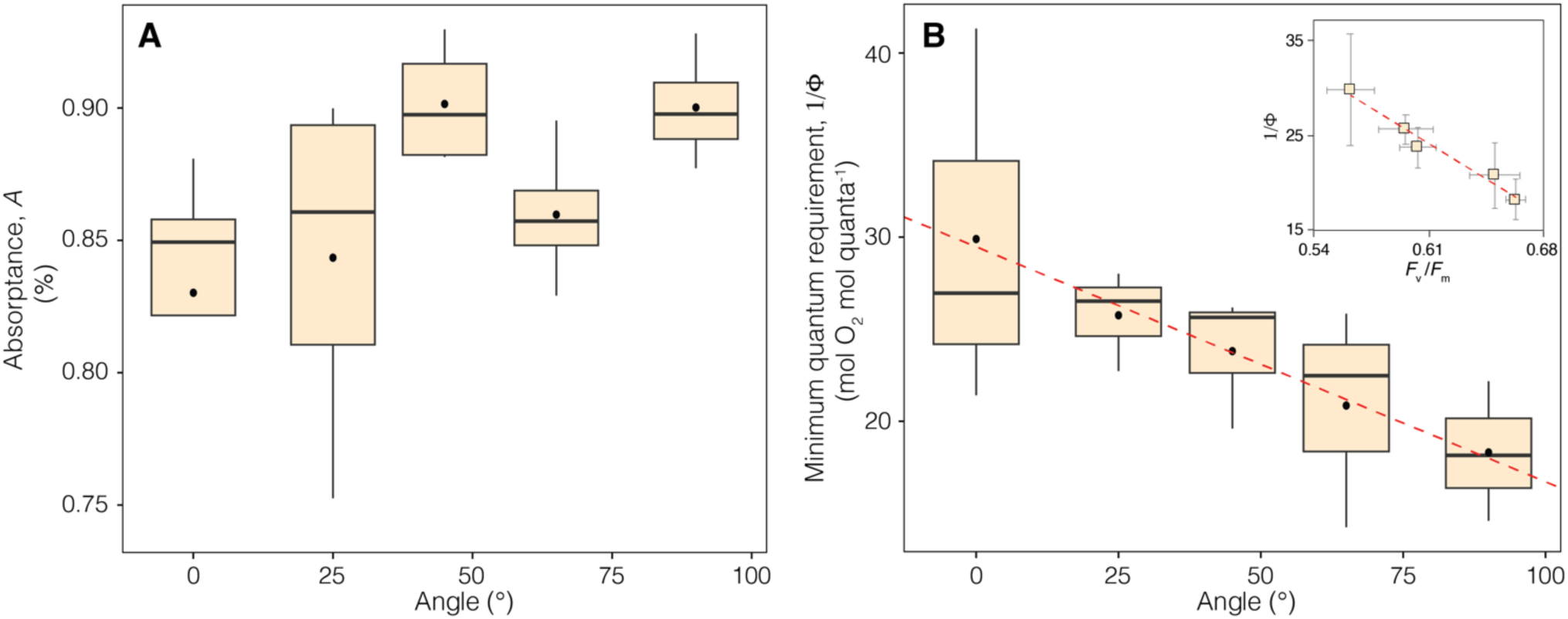
Variation in optical properties of the coral tissues and associated photosynthetic descriptors with coral surface inclination. (A) Boxplots describing the variation in coral absorptance at 675 nm (*A*675), and (B) minimum quantum requirements for oxygen evolution (1/*Φ*) with coral surface inclination. The insert in (B) depict the linear relationship between 1/*Φ* and maximum quantum yield of PSII (*F*v/*F*m), with error bars as SE.

Characterizations of the optical properties of *O. faveolata* [37] have previously shown that *A*_675_ reaches its maximum at a relatively low coral pigmentation and remains nearly constant above this threshold [29]. This explains the limited capacity of this proxy for explaining the pigment content variation in the experimental corals.

The inclination angle of the coral surface explained 40% (R^2^ = 0.40, p = 0.011) of the variation in the minimum quantum requirement for oxygen evolution (1/*Φ*), a crucial descriptor of photosynthetic performance in primary producers[31]. On average, the values of this parameter were 63% lower in vertically oriented corals exposed to the lowest irradiance (18.31 ± 3.79 quanta O_2-1_) compared to horizontally oriented corals exposed to the highest levels of direct sunlight (29.90 ± 10.27 quanta O_2-1_) (**Fig 3B**). Furthermore, a significant negative correlation was observed between the averaged values of 1/*Φ* and *F*_v_ /*F*_m_ at each inclination setting (r_(3)_ = -0.98, p = 0.002) (**Fig 3B**).

This finding indicates that the variation in 1/*Φ* along the inclination gradient was strongly associated with the photochemical energy conversion efficiency of PSII reaction centers.

### Photosynthetic parameters

Photosynthetic parameters derived from the P vs E curves were also variable in experimental corals, and a significant portion of this variation was found to be explained by their surface inclination angle. Linear regression analyses indicated that the inclination angle explained 48%, 32%, and 38% of the respective variation in the slope of the linear increase in photosynthesis at subsaturating irradiance (or photosynthetic efficiency, *α*) (R^2^ = 0.48, p < 0.01), compensating irradiance (*E*_c_) (R^2^ = 0.32, p = 0.03) and saturating irradiance (*E*_k_) (R^2^ = 0.38, p = 0.01) (**Fig 4A-C**). On average, *α, E*_c_ and *E*_k_ were respectively 42% lower, 96% higher, and 70% higher on horizontally oriented corals compared to vertically oriented ones. No significant association was observed between the variation in maximum photosynthesis (*P*_max_) and respiration rates (*R*_d_) with coral surface inclination (R^2^ = 0.01, p = 0.75 and R^2^ = 0.01, p = 0.67, respectively). However, a strong positive correlation (Pearson-product) was found between *P*_max_ and *R*_d_, regardless of coral surface inclination (r_(13)_ = 0.76, p < 0.001) (**Fig 4D**). This correlation suggests that both parameters respond similarly, although did not vary linearly along the surface inclination gradient examined in this study. The limited sample size at each inclination setting hindered a more detailed analysis of the non-linear variation observed in P_max_ and R_d_ along the experimentally induced light gradient. Furthermore, the contribution of other factors not accounted in this study cannot be discharged.

**Fig 4.**
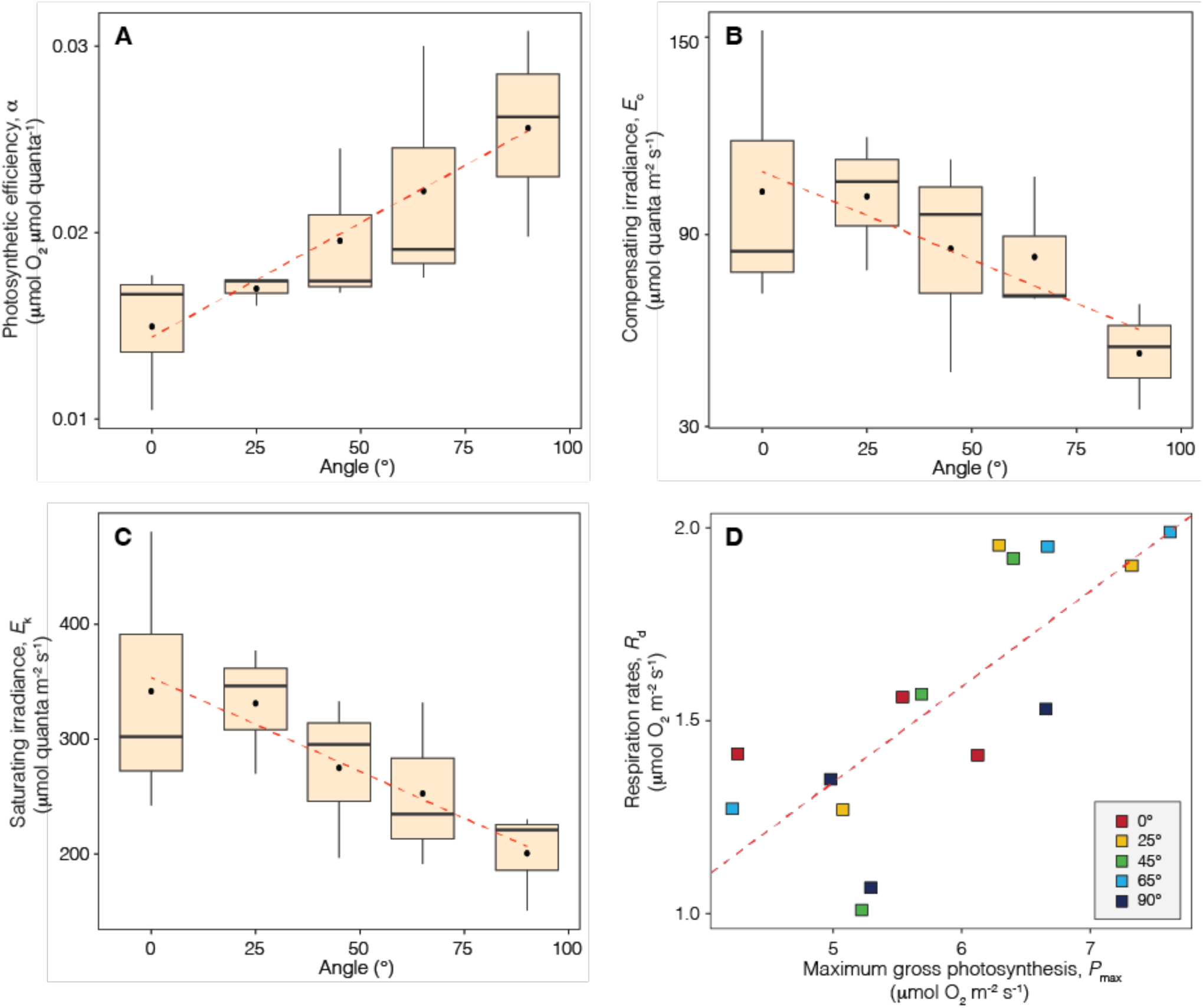
Variation in photosynthetic parameters with the inclination angle of coral surface. (A-C) Boxplots describing the variation of three photosynthetic parameters (α, *E*c and *E*k), along with linear regressions showing the association between each parameter and coral surface inclination. (D) Linear association between *P*max and *R*d of all coral samples analyzed in this study.

### Local irradiance regimes

During 143 days of irradiance data and assuming a constant photoacclimatory condition in the experimental samples, the daily light integral (DLI) received by corals in horizontal position was estimated to be 17.07 ± 4.36 mol quanta m^-2^ d^-1^, while for corals in vertical position, the DLI was reduced to 1.62 ± 0.41 mol quanta m^-2^ d^-1^. The different components of the DLI which corresponded to sub-compensating (below *E*_c_), compensating (between *E*_c_ and *E*_k_) and saturating irradiance (above *E*_k_) exhibited high variability along the inclination gradient (**Fig 5A, B**). Nearly 1/3 of the DLI experienced by horizontally oriented corals at 0° exceeded *E*_k_. This light regime was not expected to induce increases in the photosynthetic capacity but rather solely to increases in the amount of solar energy absorbed in excess by the photosynthetic apparatus of the zooxanthellae. The amount of excess solar energy absorbed gradually decreased with increasing coral surface inclination until it reached negligible values in corals at an inclination angle of 65°, where it represented less than 1% of the DLI.

**Fig 5.**
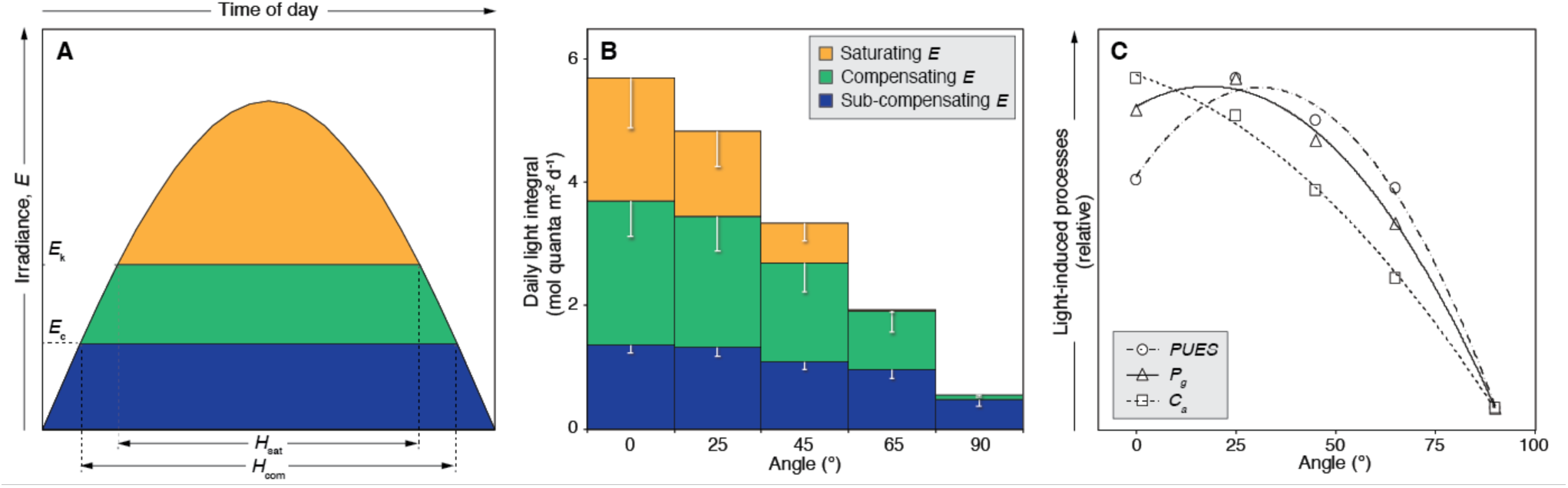
Local irradiance regimes and light-driven processes according to coral surface inclination. (A) Description of a diurnal cycle of solar irradiance (*E*), indicating the relative positions of the compensating irradiance (*E*c) and saturating irradiance (*E*k), and the daytime periods of light intensities exceeding *E*c (*H*com) and *E*k (*H*sat) [modified from [34]]. (B) Estimated daily light integral (DLI) at each inclination setting, indicating the relative proportions corresponding to sub- compensating, compensating, and saturating irradiance. Error bars describe standard deviations. (C) Estimated relative changes in light-induced processes affecting the energy balance of coral holobionts. *PUES*: photosynthetic usable energy supply; *P*g: gross productivity of the zooxanthellae; *C*a: metabolic costs of maintenance associated with zooxanthellae photosynthesis. Polynomial regressions were used to depict the relationships of these parameters with surface inclination angle.

In contrast, most of the DLI available for vertically oriented corals at 90° (90%), corresponded to sub-compensating levels of irradiance (i.e., below *E*_c_), and therefore to conditions were the respiratory demands of oxygen of the coral holobiont exceeded the photosynthetic capacity of the zooxanthellae (**Fig 5B**). Accordingly, the estimated periods of daytime when light intensity exceeded the compensating irradiance (*H*_com_) and saturating irradiance (*H*_sat_), were highly variable along the inclination gradient. On average, the *H*_com_ period for corals in horizontal position accounted for approximately 70% of the daytime (9.85 ± 1.60 h), while for vertically oriented corals it was reduced to 30% (4.16 ± 1.78 h). Notably, on some days, corals at inclination angles of 65° and 90° experienced light-limiting conditions throughout the entire day, where irradiance levels never reached *E*_c_. In contrast, the *H*_sat_ period for horizontally oriented corals comprised 48% of the daytime (6.70 ± 2.18 h), whereas for corals at 65° it was reduced to 3% (0.44 ± 0.69 h). This analysis suggests that vertically oriented corals never reached maximum photosynthetic capacity.

### Light-driven processes

The estimated photosynthetic production (*P*_g_) exhibited the highest values in corals at inclination angles of 25° (0.17 ± 0.03 mol O_2_ m^-2^ d^-1^) and 0° (0.15 ± 0.03 mol O_2_ m^-2^ d^-1^), while the lowest values were observed in corals at an inclination angle of 90° (0.04 ± 0.001 mol O_2_ m^-2^ d^-1^) (**Fig 5C**). This pattern can be explained by the hyperbolic response of photosynthesis to increasing light intensity, where no further increases in photosynthetic production occur above *E*_k_ despite higher light availability. On the other hand, the estimated energy costs of repair from photodamage for the symbiotic algae (*C*_a_), which are expected to be proportional to light availability [26], ranged between 49% and 15% of the photosynthetically fixed energy at both ends of the inclination gradient.

Consequently, the photosynthetic usable energy supplied (*PUES*) by the symbiotic algae to their coral host followed a unimodal pattern across the inclination gradient, being higher in coral surfaces at 25° and 45° inclination angles. Despite the changes in photoacclimation, the model indicated that *PUES* for fully vertically oriented corals is negligible, potentially compromising the growth and long-term survival of the experimental samples (**Fig 5C**).

## Discussion

Colony surface geometry generates, within a single depth, a diverse array of local light climates that led to significant variation in the photoacclimatory responses of the coral holobiont. Increasing the inclination angle of coral surface results in a proportional reduction of light availability, following the cosine law [16] and influenced by relative changes in the diffuse and direct components of total irradiance. While changes in light availability are commonly associated with depth gradients and the optical properties of the water column, the impact of colony surface geometry on light availability for the holobiont is often overlooked, despite the pronounced intensity gradient observed in this study. The extremes within this gradient were comparable to the estimated changes in downwelling irradiance across depths separated by approximately 40 m in clear waters (with a vertical attenuation coefficient for downwelling irradiance*, K*_d_, of 0.06 m^-1^ [38]). Consistent with previous studies [12,14,15], the results of this research provide compelling evidence that the light gradients taking place along broad depth ranges can also manifest within a single colony. The variability in these local light climates elicits comparable photoacclimation responses to optimize the balance between absorbed light energy, photosynthetic rates, photodamage and repair processes, and the translocation of photosynthetic energy by the symbiotic algae to their coral host.

The light gradient experimentally induced by changing coral surface inclination resulted in contrasting levels of PSII excitation pressure, which is one of the first chloroplastic redox signal to trigger photoacclimatory changes in primary producers [17,39]. Variation in the maximum excitation pressure over PSII (*Q*_m_) along the inclination gradient reflects the magnitude of diurnal reductions in the effective PSII quantum yields at noon (Δ*F*/*F*_m_’) relative to the maximum PSII quantum yields at dawn or dusk (*F*_v_/*F*_m_) [27]. Samples with higher inclination angles exposed to lower light intensities (at 65° and 90°), exhibited minimal diurnal oscillations in PSII quantum yields (i.e., small differences between Δ*F*/*F*_m_’ and *F*_v_/*F*_m_) and, consequently, lower values of *Q*_m_. This pattern suggests that the photosynthetic activity of the zooxanthellae at peak noon irradiance is insignificant, resulting in a potentially negligible energetic contribution to their coral host [27]. In contrast, coral samples with lower inclination angles (at 0° and 25°) exposed to higher levels of direct sunlight, exhibited substantial reductions of Δ*F*/*F*_m_’ at noon relative to *F*_v_/*F*_m_ at dusk/dawn, leading to higher values of *Q*_m_. This indicates that during peak-irradiance periods, a significant portion of PSII reaction centers becomes closed due to the high rate of energy absorbed (light dose), requiring maximum induction of photoprotective mechanisms to dissipate as heat the excess absorbed energy [27,40,41]. Accordingly, the variation in *F*_v_/*F*_m_ along the inclination gradient reflects a variable proportion of photoinactive PSII reaction centers that are not involved in photochemistry but are better suited to dissipate excess excitation energy [42,43]. The observed linear association between *F*_v_/*F*_m_ and surface inclination angle suggests that the subpopulation of photoinactive PSII in the algal photosynthetic apparatus may be inversely proportional to the inclination angle of coral surface, resulting in maximum photoprotection on horizontal surfaces and maximal photochemical efficiency on vertical surfaces.

The consistency between the patterns obtained in experimental samples and in coral colonies within natural environments, suggests that the observed changes in PSII photochemical efficiency induced by coral adjustments to surface inclination also occur naturally in coral colonies with complex geometries [39,43].

The induction of long-term photoacclimatory responses in coral samples was supported by the observed variation in their *in vivo* optical properties (absorptance/pigmentation) and photosynthetic potential (P vs E curve parameters). Although the relationship between light absorption capacity (*A*_675_) and coral surface inclination angle was weak, it still indicated a reduction in pigment content in coral samples with lower inclination angles that were exposed to higher levels of direct sunlight, which exhibited higher PSII excitation pressure. Such inferred changes in pigmentation along the inclination gradient are consistent with the induction of a photoacclimatory response to reduce excitation pressure on PSII in high light environments and to maximize the light harvesting capacity in low light conditions. Coral samples exposed to higher inclinations (above 45°) displayed similar *A*_675_ values close to 0.9, which is the maximum light absorption capacity at 675 nm documented for *O. faveolata* [37]. This suggests that the light absorption capacity of coral holobionts was already close to its maximum and minimally influenced by further changes in pigment content. These findings highlight the importance of absorptance variation associated with changes in pigmentation as a crucial photoacclimatory response to modulate light dose, not only for coral colonies located at different depths [19,20,22] but also for coral surfaces within a single colony exhibiting contrasting inclination.

Comparative analyses of the P vs E curve parameters revealed clear linear associations between the photosynthetic efficiency (*α*), compensating irradiance (*E*_c_), and saturating irradiance (*E*_k_) with the inclination angle of the coral surface. The pattern of variation in these parameters is similar to that observed in corals growing under contrasting light climates along depth gradients [19,20,22], being directly associated with the observed changes in absorptance/pigmentation and quantum yield efficiency of PSII. Several outcomes result from the association between these parameters and coral surface inclination. Firstly, the enhanced light utilization efficiency in coral surfaces with greater inclination angles leads to more rapid increases in photosynthetic rates with increasing light -availability. Secondly, the lowered compensating irradiance in coral surfaces with greater inclination angles and limited light availability enhances their capacity to achieve positive energy balances over extended periods of the daytime. Lastly, the elevated saturating irradiance in horizontal or slightly inclined coral surfaces exposed to higher levels of direct sunlight leads to reductions in the proportion of daytime when irradiance exceeds the maximum photosynthetic capacity, minimizing the accumulation of excessive energy absorbed that cannot be used in photosynthesis and must be dissipated through photoprotective mechanisms. Both *E*_c_ and *E*_k_ are very sensitive to changes in the photosynthetic efficiency (*E*_c_ = *R*_d_/*α* and *E*_k_ = *P*_max_/*α*). Therefore, variations in these photosynthetic parameters along the inclination gradient primarily are likely attributed to changes in the absorption cross-section of the coral surface and/or the number of PSII reaction centers in the photosynthetic apparatus of the zooxanthellae, which directly influence light utilization efficiency [17,44,45]. The analysis of the interaction between the photosynthetic parameters and the optical properties of the holobiont (the coral animal + symbiotic algae) revealed a clear pattern of change in the minimum quantum requirements of photosynthesis (1/*Φ*), measured as oxygen evolution, with coral surface inclination. The values of this descriptor of photosynthetic performance decreased proportionally with increasing coral surface inclination and decreasing irradiance, resembling the pattern of change observed in several coral species with increasing depth [20,46]. This trend indicates that coral surfaces with high inclination angles and reduced light availability exhibit higher quantum efficiency of photosynthesis compared to coral surfaces with low inclination angles and maximum light exposure. Furthermore, the correlation between 1/*Φ* and *F*_v_/*F*_m_ reinforces the fundamental role of the accumulation of photodamaged PSII reaction centers associated with light exposure in influencing the photosynthetic quantum efficiency of corals.

Analyses of the diurnal variation in irradiance along the inclination gradient revealed contrasting effects on the photosynthetic potential and energetic performance of coral holobionts. Coral surfaces with low inclination angle, directly exposed to downwelling irradiance, received sufficient light to reach maximum photosynthesis during nearly half of the daytime. The estimated photosynthetic productivity of these coral surfaces remained relatively constant regardless of slight changes in the inclination angle and light exposure. This plateau in photosynthetic productivity occurs after reaching maximum photosynthetic capacity determined by the hyperbolic response of photosynthesis to light availability, where further increases in productivity are not observed once photosynthesis is light- saturated [47]. As a result, a considerable proportion of the daily light dose absorbed by horizontal and nearly horizontal coral surfaces cannot be utilized for photosynthetic energy conversion. Instead, it represents excess excitation energy that can be harmful for the photosynthetic apparatus of the zooxanthellae [26,39,42]. This energy absorbed in excess must be dissipated through photoprotective mechanisms, a condition confirmed by the high *Q*_m_ values determined from chlorophyll *a* fluorescence measurements, which are indicative of high levels of non-photochemical quenching induced. Conversely, coral surfaces with higher inclination angles and reduced light exposure may only receive enough light the exceed the compensating irradiance during brief periods of the day and may never reach their maximum photosynthetic capacity. Our analyses suggest that coral surfaces with greater inclination angles can sustain positive energy balances only over short time periods when the photosynthetically fixed carbon exceeds the holobiont respiratory requirements. This is consistent with the low *Q*_m_ values determined for these corals, reflecting minimum photosynthetic activity of the symbiotic algae and, consequently, negligible energetic contribution to the coral host metabolism. In these coral surfaces calcification and growth capacities may be minimal, imposing a potential tolerance limit on the symbiosis.

The non-linearity between light availability, photosynthetic production and costs of repair from photodamage can result in contrasting levels of photosynthetic usable energy supplied by the zooxanthellae to their host (*PUES*), depending on the geometry of coral colonies. While photosynthetic production follows a hyperbolic response to light intensity, the amounts of absorbed light energy and the costs of repair from photodamage increase proportionally with light availability [26,47]. This energetic imbalance explains the predicted decrease in *PUES* in both vertically and horizontally oriented coral surfaces, similar to changes along depth gradients [26]. The predicted pattern of change in *PUES* as a function of the inclination angle exhibits a unimodal shape, mediated by two contrasting processes: 1) in horizontally oriented surfaces, exposure to intense irradiance amplifies the metabolic costs of maintenance in the zooxanthellae, limiting the amount of energy that can be translocated to their host; and 2) as the inclination angle increases, the reductions in light availability and photosynthetic activity lead to decreases in the amount of energy fixed in photosynthesis. In the shallow experimental site of our study, corals can achieve maximum energetic output from the zooxanthellae and energetic performance at intermediate surface inclination angles. However, with increasing depth, the optimal inclination angles for energetic performance tend to approach the horizontal orientation due to the reduction in excess light energy absorbed relative to the maximum photosynthetic capacity. This predicted trend aligns with the prevalence of flattened morphologies observed in deep, low light environments, as these morphologies allow coral colonies to maximize light capture and minimize self-shading [48,49].

Overall, the combination of water optical properties [13,50], substrate type and landscape architecture [51,52], determines the balance between direct and diffuse components in the local light field at specific depths. The vertical attenuation coefficient for downwelling irradiance (*K*_d_), a proxy of the water optical properties, is influenced by the concentration and composition of dissolved and particulate materials, which can absorb and scatter light as it travels through the water. As light penetrates deeper into the water column, there is an increased likelihood of scattering, resulting in an increased proportion of the diffuse component (i.e., the light coming from aside) with depth [16].

This effect may favor light interception and energy acquisition in non-horizontal surfaces of coral colonies located at intermediate depths, where light is still not limited [13,38]. Additionally, different substrates have varying reflectance and absorption properties, further influencing the light regime experienced by benthic photosynthetic organisms (e.g., amplitude, intensity and symmetry), and thus their primary productivity [51,52]. We acknowledge that all these factors along with others, can contribute to the complexity of underwater light behavior [53]. However, the framework of this study specifically focuses on investigating the effect of variations in light availability associated with surface inclination on the photoacclimatory response of coral holobionts, minimizing the potential influence of other biotic and abiotic conditions that can also modulate colony light availability.

In our study, we used small coral samples detached from the original colony. However, it is important to note that under natural conditions, coral colonies with complex morphologies are physiologically integrated, enabling the translocation of energetic products from source to sink sites, such as growing tips and regenerating areas [5–7,54]. This translocation allows for the highest calcification rates in tips of branching corals, which typically have low zooxanthellae densities and rates of net primary productivity [5,7]. Similarly, in massive corals, the growth of colony margins and tissue regeneration is promoted by the translocation of photosynthetic products from the most productive areas of the colony [6,54]. Although our study focused on individual samples and did not directly assess within-colony translocation, the evidence from previous studies indicates that the intracolonial translocation of photosynthetically derived energy products may be a common feature among colonial corals, occurring from highly productive source sites to regions with higher metabolic demands. Our study findings align with the evidence that coral colonies with complex geometries exhibit great heterogeneity in terms of local productivity, energetic demands and energy fluxes. This supports the importance of colony integration and the translocation of energetic products in enhancing the overall performance of the entire coral colony.

In shallow, high-light environments, horizontal coral surfaces, which are typically located in the central areas of colonies, may not coincide with the most productive regions of the colony as a result of the increased energy costs of repair from photodamage in the zooxanthellae, which ultimately affects the amount of photosynthetic usable energy supplied to their coral host (*PUES*) [26]. The greater extension and calcification commonly observed in these areas [55] potentially result from energetic products imported from other more productive source sites within the colony. On the other hand, coral surfaces with a large inclination angle and reduced productivity may also act as energy sinks within the colony. Survival of these surfaces would require energetic subsidies from more productive areas of the colony due to their extremely low levels of *PUES*. As depth increases, these low-productivity areas with greater inclination angles are predicted to experience negative photosynthetic energy balances constantly, becoming increasingly less productive and more costly to maintain for the colony. The reduced productivity of colony surfaces with greater inclination angles, along with their partial dependence on other parts of the colony for energy, may partially explain their higher vulnerability to reduced light penetration associated with increased water turbidity [50] (**Fig 6A**), disease outbreaks such as the recent stony coral tissue loss disease [56], as well as environmental perturbations, including heat stress [57,58] (**Fig 6B**).

**Fig 6.**
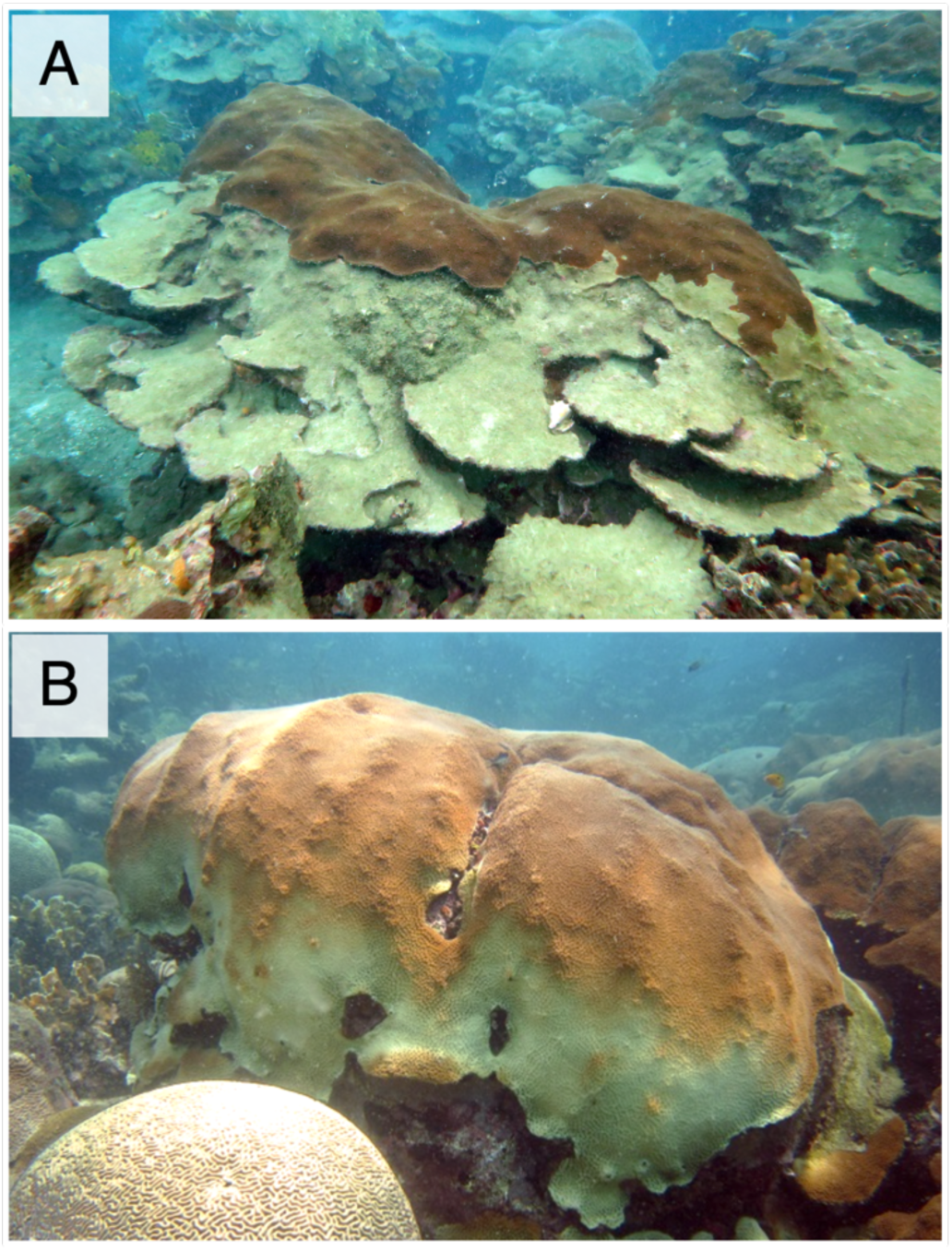
Coral colonies of *O. faveolata* affected by environmental perturbations. (A) Partial mortality observed in a coral colony in the Varadero Reef, an area experiencing elevated anthropogenic turbidity [50]. (B) Bleached and non-bleached regions within a colony during the widespread bleaching event that occurred in the Caribbean in 2010. It is noteworthy that in both conditions, the most affected areas are the low-productivity colony surfaces with the larger inclination angles.

In conclusion, this study highlights the significant influence of colony geometry on light availability, holobiont physiology and photosynthetic energy acquisition in zooxanthellate corals. Previous research has shown that variations in photoacclimation responses of coral holobionts can be linked to changes in the composition of symbiotic zooxanthellae along environmental gradients [21,27,58,59]. Most corals with broad bathymetric distributions, including *O. faveolata*, exhibit changes in the symbiont community composition which presumably allow them to photoacclimate to particular light environments, both across depths and within colonies [58,60]. Thus, part of the observed variation in holobiont physiological parameters, both within and between inclination settings, can potentially be attributed to changes in *Symbiodiniaceae* community composition or other factors related to particular metabolic conditions of experimental corals. However, a significant portion of this variation can be solely attributed to the inclination angle of the coral surface and the associated light climate experimentally induced in this study. This indicates that light energy availability remains a major driver of variation in coral holobiont physiology. Our study provides clear evidence that contrasting photoacclimatory responses, light climates and energy balances can occur within a single colony at a constant depth. Therefore, studying the entire coral colony and its metabolic gradients should be a research priority to enhance our understanding of colonial responses to environmental cues and the role of colony morphology and energetic economy on the vertical distribution of species and niche diversification in coral reefs.

## Acknowledgments

Claudia T. Galindo-Martínez, Kelly Gómez-Campo, Luis A. González-Guerrero, Ricardo I. Cruz-Cano, Nancy Escandón-Flores, Nadine Schubert and Darren Brown provided support during field activities and laboratory analysis. The National Commission on Aquaculture and Fisheries (CONAPESCA) for research permit (DGOPA 08606.251011.3021). We also thank the DGAPA-UNAM for the financial support of a sabbatical period at PSU to SE with a PASPA fellowship.

## Notes

https://figshare.com/s/1acd845747e8ba3d3d92

### Competing Interest Statement

The authors have declared no competing interest.

